# Mapping endothelial functional phenotype in cancer by unveiling the kinase and phosphatase drivers

**DOI:** 10.1101/2020.07.14.201988

**Authors:** Or Gadish, Elazer R. Edelman

## Abstract

Endothelial cells (EC) are state-dependent regulators of the tumor ecosystem: quiescent ECs promote homeostasis; proliferative ECs stimulate tumor growth. Tumors, in turn, promote pro-tumorigenic EC phenotype. We studied functional and phosphorylative transformations on EC state in cancer. Quiescent HUVECs cultured in breast cancer cell-conditioned media displayed marked elongation and impaired wound healing. Quantitative mass spectrometry identified phosphorylative regulators of this dysfunctional transformation. Growth factor receptor kinases showed decreased, rather than increased activity, suggesting that EC regulation in tumors can arise other than from classic growth-factor-mediated angiogenesis alone. Of the 152 kinases and phosphatases across 62 families, six were chosen for functional validation using pharmacologic inhibitors. Inhibiting Akt and Ptp1b restored EC regulatory state, warranting further investigation as therapeutic targets; Src inhibition, however, promoted the dysfunctional phenotype, suggesting caution for Src inhibitors as EC-regulating therapies. Mapping phosphorylative drivers reveals complex relationships between EC phenotype, transformation, and regulation, and may shed light on how existing cancer-targeting inhibitors affect tumor endothelium. Data are available via ProteomeXchange with identifier PXD020333.

## Introduction

Endothelial cell (EC) state is critical to the regulation of the surrounding tissue: quiescent ECs promote homeostasis, while proliferative ECs stimulate growth (Nugent *et al*, 1993; Ettenson *et al*, 2000). This aspect of EC biology holds for cancer biology as it does for processes like hyperplasia, protein infiltration and infection: quiescent ECs promote homeostasis, while proliferative ECs stimulate tumor growth (Butler *et al*, 2010; Ferraro *et al*, 2019; Ghiabi *et al*, 2015). There seems to be a particularly powerful dynamic between tumors and ECs wherein cancer cells secrete factors and create physical cancer-EC conduits that promote emergence of pro-tumorigenic phenotype in ECs (Folkman, 1971; Senger *et al*, 1983; Butler *et al*, 2010; Zhou *et al*, 2014; Ghiabi *et al*, 2015; Ferraro *et al*, 2019; Connor *et al*, 2015). Normalization of EC state as an adjuvant to existing therapies has the potential to revive anti-tumor immune response and reverse overall tumor progression (Jain, 2005; Martin *et al*, 2019).

The intact endothelial cell (EC) state and phenotype create a robust environment for optimized tissue repair. Quiescent ECs have maximal regulatory capacity (Nugent *et al*, 1993; Ettenson *et al*, 2000); whereas proliferative ECs (such as those involved in angiogenesis) can have opposite, dysfunctional and cancer promoting effects (Ferraro *et al*, 2019; Butler *et al*, 2010; Ghiabi *et al*, 2015).

Current methods of tumor vascular normalization have generated much enthusiasm, but are not the only avenue for leveraging endothelial biology in cancer (Jain, 2008; Giuliano & Pagès, 2013; Martin *et al*, 2019). Recently, the response of confluent, quiescent ECs to breast-cancer-conditioned media was shown to be *inversely* proportional to levels of VEGF, a major target of tradition vascular normalization (Gomez *et al*, 2016). In fact, conditioned media from more aggressive cancer cell lines had lower VEGF; media with lower VEGF, however, resulted in increased EC elongation. Since state affects EC response to tumors, these data suggest that, in addition to classic GF-mediated angiogenesis, there are other means of EC regulation in cancer and demand that we consider non-growth factor biology.

Cytoskeletal regulation is a critical determinant and aspect of EC state (Indolfi *et al*, 2012; Gomez *et al*, 2016), and weaker cell-substrate interactions in tumor ECs result not only in increased permeability, but also proliferation and migration (Bogatcheva & Verin, 2008; Xiong *et al*, 2009). EC cytoskeletal reorganization, like many processes in mammalian cells, is regulated in large part by phosphorylation and phosphorylative modifications pervade as much as 90% of the human proteome (Sharma *et al*, 2014). Yet, phosphorylative regulation has been studied significantly less than RNA expression or protein translation in cancer. The main challenges are: quantification of changes in phosphorylation that occur on the order of seconds to minutes (Olsen *et al*, 2006); identifying the specific effects of phosphorylation (activating, inhibitory, or with no effect), which depend on the particular site and protein (Sharma *et al*, 2014); and difficulty of high-throughput analysis. Recent advances in phosphopeptide detection by mass spectrometry (Sharma *et al*, 2014; Olsen *et al*, 2006, 2010), circumvents the technical limitations and paves the way for more in-depth investigation.

Libraries detailing the effects of phosphosites on each protein, such as PhosphoSitePlus (Hornbeck *et al*, 2012), can aid in determining the implications of phosphorylative changes; however, these libraries are still too sparse to analyze overall patterns in phosphoproteomic datasets. An alternate approach is to use changes in phosphorylation as a proxy for the activity of the kinases and phosphatases that catalyze the addition or removal of phosphates at that phosphosite (hereafter jointly referred to as “phosphoenzymes”). Indeed, phosphoenzymes make excellent therapeutic targets and kinase inhibitors are the current state of the art for targeting tumor vasculature, with eleven recently approved or in phase II/III clinical trials (Qin *et al*, 2019).

This study specifically explored the transformation of ECs by triple-negative breast cancer (TNBC), the subtype of breast cancer with highest recurrence and mortality, and worst response to chemotherapy (Reddy *et al*, 2018). TNBC is furthermore, an ideal model of study *in vitro* as lacking hormone receptors, these cells are relatively isolated from endocrinological control *in vivo*, increasing the validity of biological results *in vitro*. In contrast to previous studies, we examined the phosphorylative regulation in ECs grown in cancer-conditioned media and focused on ECs that began in their quiescent, growth-inhibitory state, rather than the promoting, exponential growth phase.

## Results

### Breast-cancer-conditioned media impairs wound healing in concert with morphologic changes in endothelial cells

Unlike many tumor EC models which examine angiogenesis during exponential EC growth, ECs in our model were first grown to confluence—and into what has been termed phenotypic *quiescence*—before exposure to cancer-conditioned media (CCM) or fresh growth media (CTRL). CCM was collected for 2 days from triple-negative MDA-MB-231 breast cancer cells after 4 days in culture, and then transferred to the confluent human umbilical vein endothelial cell cultures (HUVEC). ECs in CCM exhibit robustly increased elongation after 36-48 hours of co-culture, clearly visible by phase contrast microscopy (Figure 1A-B). The formation of swirl patterns suggests a locally heterogeneous response, and possible sensitivity to local flow domains.

**Figure 1.**
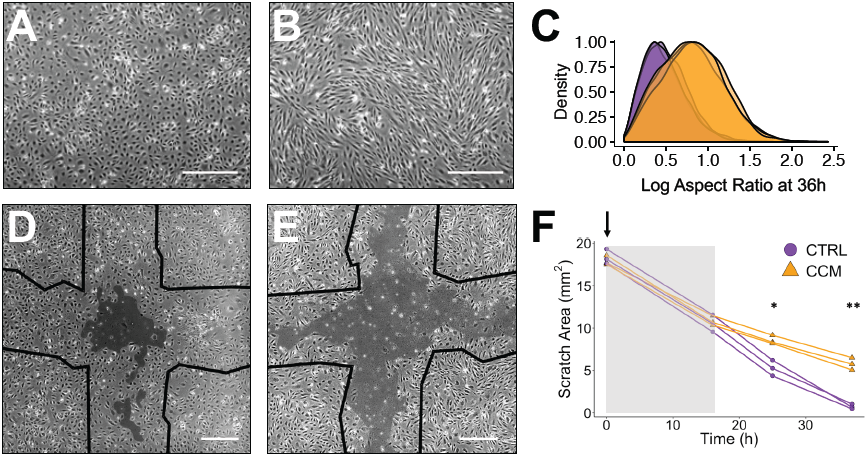
Breast cancer-conditioned media drives EC morphologic transformation and wound healing. Representative phase contrast micrographs of HUVEC culture show variable elongation after 36 hours in control (A, CTRL) or cancer-conditioned media (B, CCM). Kernel-estimated density plots of elongation (log aspect ratio) across replicates quantitatively separate phenotype (C). Representative micrographs of the unclosed HUVEC area at the center of a cross-scratch assay after 36 hours similarly differ in CTRL (D) or CCM (E). For the first 16 hours after start of conditioning (F, arrow, shaded area), there is no significant difference between scratch area, but afterwards the pattern diverges (F). *Scale bar: 200 µm. *p < 0*.*05, **p < 0*.*01, CCM vs. CTRL*.

To quantify morphology, fluorescent cell-filling Cell Tracker dyes were used with semi-automatic segmentation by ImageJ. We were specifically intent on quantifying elongation and sought an elongation metric that was normally distributed among our cell populations and maximized separation between CTRL and CCM distributions. The Log Aspect Ratio (LAR)—the natural logarithm of the aspect ratio of an elliptical fit—was the ideal choice: it is unbounded in the direction from CTRL to CCM and is pseudo-normally distributed (Expanded View Figure 1). LAR quantifiably reproduces the visual difference: ECs in CCM showed a 77% increase in median LAR from CTRL (0.455 to 0.807), and a 118% increase in the mode (Figure 1C). Other common metrics, such as circularity, eccentricity, and the non-normalized aspect ratio, were considered but were inferior in separation and/or distribution (Expanded View Figure 1).

To map functional phenotype, we probed EC cytoskeletal transformations under stress of controlled scratch injury to the EC monolayer. ECs in CCM displayed markedly impaired monolayer reconstruction (Figure 1D-F). By the end of 36 hours, the remaining acellular scratch area was 6.8-fold smaller in CTRL than in CCM (Figure 1F). Control ECs closed 96.1 ± 1.6% of the scratch area by 36 hours, but ECs in CCM only 67.8 ± 4.1%. Furthermore, the reduced closure rate started at ∼16 hours after CCM exposure, matching the progression of morphologic transformation and strongly suggesting that wound closure represents the same underlying cytoskeletal transformation as elongation.

Importantly, all CCM conditions used a 3:1 ratio of CCM to CTRL media and supplemented with glucose and bicarbonate to offset possible nutrient depletion. Bicarbonate supplementation was especially important given the low levels in normal human endothelial growth media, which causes increased lactic acid in conditioned media. Depletion of other nutrients was difficult to ascertain, but experiments using CCM concentrated by centrifugal filtration showed similar morphologic changes.

### Endothelial phosphoproteomic profile shifts significantly in breast cancer conditioned media

Cytoskeletal reorganization in ECs is regulated in large part by phosphorylation. We used mass spectrometry-based phosphoproteomics on HUVEC lysates to quantify the levels of 2,602 nonredundant phosphorylated peptides representing 1,243 distinct proteins (Figure 2A). Overall, the phosphoproteomic profiles of ECs in CTRL and CCM are statistically distinct (p < 0.005 by clustering analysis, Figure 2A).

**Figure 2.**
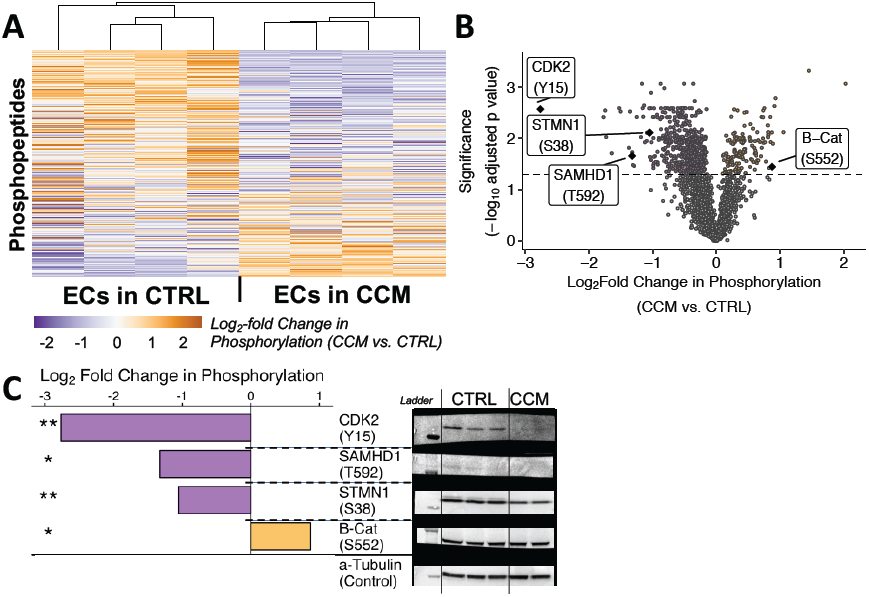
Phosphoproteomic profile of ECs shifts with cancer conditioning. High-throughput phosphoproteomics of 2,602 phosphosites revealed two distinct phenotypes between CTRL and CCM (A), and yielded 668 significantly regulated sites (B), four of which were separately validated by Western blotting (C).

The chosen phosphoproteomic workflow included a phosphotyrosine (pTyr) enrichment step in addition to the typical phosphoenrichment achieved by metal oxide affinity chromatography. Phosphotyrosines are extremely important for cellular activity, even though they account for only 0.4% of all phosphosites (Sharma *et al*, 2014). This enrichment increased our pTyr coverage to 4.2%.

We found 668 phosphorylation sites with statistically different levels between CTRL and CCM, referred to hereafter as “regulated sites” (Figure 2B). Consistent with previous studies reporting increased pTyr proportion among regulated sites (Olsen *et al*, 2006), pTyr proportion increased in our data from 4.2% of all sites to 5.8% of regulated sites.

Four regulated sites were chosen for subsequent analysis by Western Blot (Figure 2C) from among those with the largest absolute change and for which high quality antibodies were readily available. Downregulation in CCM was validated for pTyr15 on cyclin-dependent kinase 1/2 (CDK2_Y15); pSer38 on Stathmin 1 (STMN1_S38); pThr592 on SAM domain- and HD domain-containing protein 1 (SAMHD1_T592), despite markedly reduced overall staining in the case of SAMHD1. CCM upregulation of pSer552 on beta Catenin (B-Cat_S552) was not reproduced by Western blot, despite robust overall staining. This reminder of how high-throughput analysis may amplify statistical anomalies led us to seek macropatterns to map underlying cellular activity, rather than individual differentially regulated sites.

### Cancer-conditioned phosphoproteomic shift is associated with cytoskeletal regulation

We then tested whether phosphoproteomic variation mapped to cytoskeletal changes. Gene set overlap analysis (Subramanian *et al*, 2005; Liberzon *et al*, 2011) confirmed that cytoskeletal gene sets were enriched overall among our differentially regulated sites. From the Gene Ontology Cellular Component (GO CC) collection (Ashburner *et al*, 2000; The Gene Ontology Consortium, 2019), 9 of the top 10 most significant gene sets were involved in the regulation of cytoskeletal components, topped by *GO_CYTOSKELETON* (Figure 3A). Several other prominent collections—including GO Biological Processes (GO BP) (Ashburner *et al*, 2000; The Gene Ontology Consortium, 2019), BioCarta (Nishimura, 2002) and KEGG (Kanehisa & Goto, 2000; Kanehisa *et al*, 2017, 2019)— also had significant enrichment by cytoskeletal gene sets (Expanded View Figure 2A).

**Figure 3.**
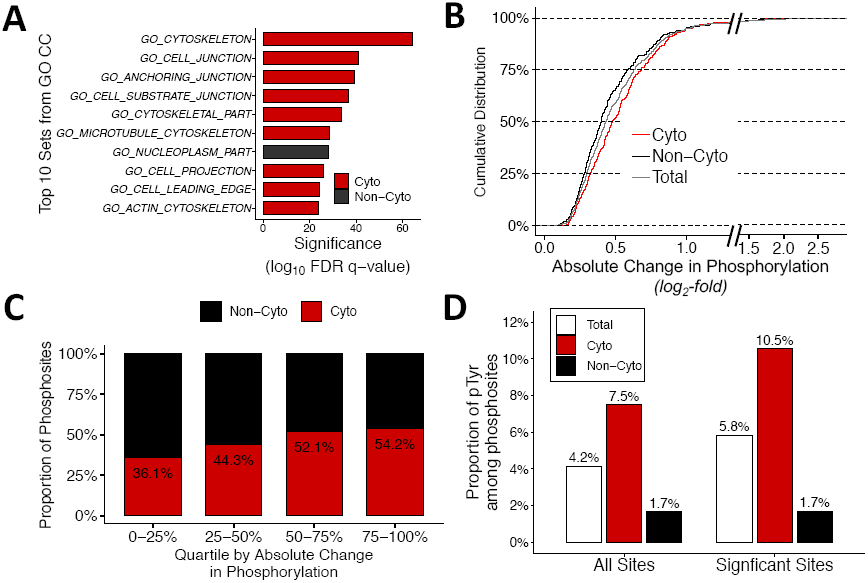
Phosphoproteomic data is associated with cytoskeletal changes. Based on gene set overlap analysis with GO Cellular Component (GO CC) collection, 9 of the top 10 gene sets are cytoskeletal (A). There is a demonstrated positive association between cytoskeletal annotation (Cyto vs Non-Cyto) and absolute change in phosphorylation, both by overall distribution (B) and when tallying sites from Cyto proteins. The proportion of phospho-tyrosine (pTyr) phosphosites is increased in Cyto proteins, when considering all sites, as well as only the significantly regulated sites (D).

Furthermore, 313 cytoskeleton-associated regulated phosphosites correlated with larger phosphorylative changes from CTRL to CCM. Additionally, cytoskeletal proteins had 26% more phosphorylation sites per protein on average compared to non-cytoskeletal proteins (p < 0.05, Expanded View Figure 2B). Classification of cytoskeletal proteins was performed using keywords from the GO CC and GO BP annotations (Ashburner *et al*, 2000; The Gene Ontology Consortium, 2019) (see Methods). Cytoskeletal sites had a 19.8% higher median change in phosphorylation than non-cytoskeletal sites (p < 0.005 by Kolmogorov-Smirnov test, Figure 3C). Absolute, log_2_-fold change was quantified, without directionality, since magnitude of change in phosphorylation correlates with activity of upstream phosphorylative enzymes. Furthermore, by tallying the proportion of cytoskeletal sites within magnitude-of-change quartiles, a similar pattern emerges: cytoskeletal proportion increasing with magnitude of change (Figure 3D). Finally, the increased proportion of pTyr peptides amongst cytoskeletal sites (Figure 3B) further establishes the role of these sites in the CCM-induced changes.

### Downstream phosphoproteomic changes predict activity of upstream kinases and phosphatases

Direct mapping to protein activity is hampered by limited availability of phosphosite annotation; instead, phosphorylative regulation was estimated by analyzing the predicted activity of upstream kinases and phosphatases (“phosphoenzymes”). The NetworKIN tool on the KinomeXplorer platform (Horn *et al*, 2014) was used to predict the phosphoenzymes directly upstream of the 313 cytoskeleton-associated regulated phosphosites. KinomeXplorer predictions employ protein network proximity analysis together with amino acid motif analysis (Figure 4A). NetworKIN scores greater than or equal to 2 (the default) were considered significant. In the specific case of phosphatases—for which the platform has limited coverage—predictions were considered significant if no other enzyme received a higher score for that phosphosite, even if the NetworKIN score was below 2. This analysis yielded 108 and 16 unique kinases and phosphatases, respectively, representing 55 phosphoenzyme families.

**Figure 4.**
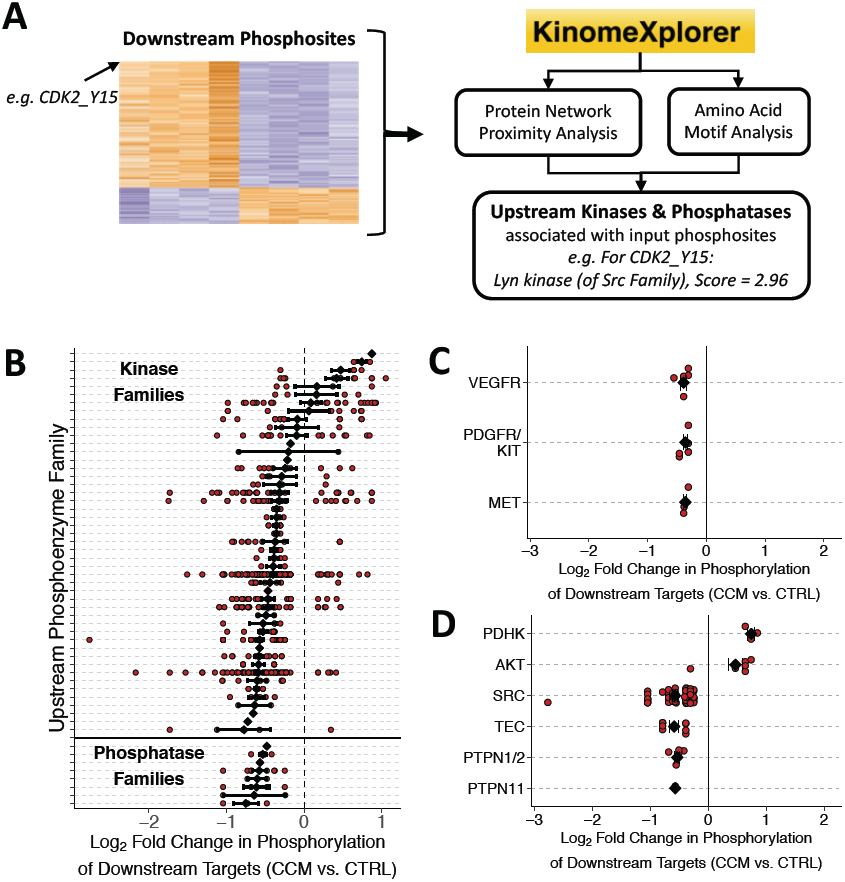
KinomeXplorer analysis maps phosphoenzymes families to quantitative phosphoproteomic data. KinomeXplorer analysis combines protein network analysis and amino acid motif analysis to predict the enzymes upstream of the significant phosphosites provided by our phosphoproteomic analysis (A). Grouped by phosphoenzyme family, activity can be estimated for all kinase and phosphatase families by mapping the change in phosphorylation of downstream targets (B). The cytoskeletal targets of four angiogenic growth factor receptor families all suggest decreased activity in CCM (C). Six families with the most differentially regulated cytoskeletal targets were chosen for further evaluation (D).

The NetworKIN score does not represent effect size; instead, we used the phosphorylation levels of downstream sites: a kinase whose predicted targets showed increased phosphorylation in CCM was estimated to be upregulated in CCM (with increased activity). Phosphatase activity, on the other hand, is inversely proportional to target phosphorylation, since phosphatases cause dephosphorylation: a phosphatase would be upregulated if the phosphorylation of its targets is decreased.

Phosphoenzymes were analyzed by family—as defined by KinomeXplorer, *e*.*g. Src kinase family* (Figure 4B)—to increase robustness to error and account for the family-level specificity of many phosphoenzyme inhibitors. Significantly regulated phosphoenzyme families were those whose phosphorylative changes from CTRL to CCM were significantly different than zero. Due to the limited coverage of phosphatases, phosphatase families with one target with a log_2_-fold change in phosphorylation more extreme than ±0.5 were also considered significant.

Overall, significant activity satisfied three statistical filters: 1) differential phosphorylation for each phosphosite (“regulated sites”); 2) significant KinomeXplorer prediction; and 3) phosphoenzyme families with significant phosphorylative changes (“regulated families”). From this analysis emerged 25 “regulated families”: 19 kinase families downregulated in CCM, three upregulated kinase families, and three upregulated phosphatase families. The families upregulated in CCM are potential targets for normalization by inhibition.

Of immediate interest were the growth factor receptor (GFR) kinases most commonly targeted by existing anti-angiogenesis kinase inhibitor therapies: vascular endothelial GFR (VEGFR), platelet-derived GFR (PDGFR), c-Kit (receptor for stem cell factor), and c-Met (receptor for hepatocyte growth factor) (Qin *et al*, 2019). All three families were significantly *downregulated* in CCM in our data (p < 0.01, p < 0.001, p < 0.05, respectively) (Figure 4C), even though levels of soluble VEGF in the media were 18-fold *higher* in CCM on average (35.4 ± 11.7 ng/mL vs 1.97 ± 0.06 ng/mL by ELISA).

Instead of looking specifically at the GFR kinases, six other regulated families were chosen for further evaluation based on the magnitude of the estimated change in activity, documented significance to tumor EC biology, and the availability of inhibitors: PDHK, AKT, SRC, TEC, PTPN1/2, and PTPN11 (Figure 4D).

### Probing with inhibitors in morphology and wound healing assays reveals phosphoenzyme activity

Six inhibitors were chosen to probe the functional effect of the six regulated families of interest: an inhibitor against pyruvate dehydrogenase kinase 1 (Pdhk1), an Akt1/2/3 inhibitor, an inhibitor against bone marrow tyrosine kinase on chromosome X (Bmx, of the TEC family), a SRC family kinase inhibitor, an inhibitor against protein tyrosine phosphatase 1b (Ptp1b, also known as PTPN1), and a broader spectrum phosphatase inhibitor which targets Ptp1b and the SH2 domain-containing phosphatase 2 (Shp2, also known as PTPN11) (Expanded View Figure 3A). Each inhibitor was tested for HUVEC cytotoxicity at doses up to 1,000-times the reported IC50 or K_i_, and the highest non-cytotoxic dose was chosen for subsequent experiments. The statistical analysis above for determining regulated families was repeated for each inhibitor and all of its targets, and none of the secondary targets were found to affect the previous analysis (p < 0.01, Expanded View Figure 3B).

The inhibitors were added to confluent EC cultures concurrent with CCM or CTRL exposure and assessed after 36 hours for morphology and wound healing as described above. The activity of kinases and phosphatases was then determined by the effects on EC phenotype, compared to media without inhibitors (Figure 5B-C). Phenotypic reversion from CCM towards CTRL suggests *increased* activity of that phosphoenzyme in CCM. Conversely, an inhibitor driving dysfunctional phenotype from CTRL towards CCM suggests *decreased* activity in CCM. In this way, phosphoenzyme activity as predicted by phosphoproteomics could be compared to that determined by inhibition testing.

**Figure 5.**
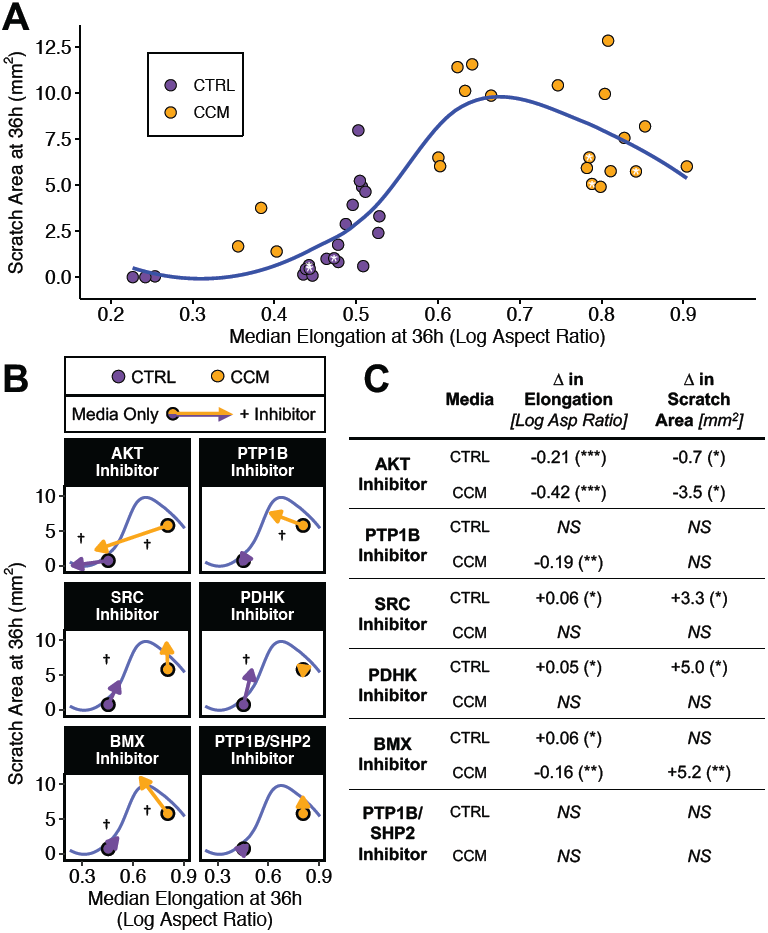
Phosphoenzyme inhibitors differentially affect morphology and wound healing. Six pharmacologic inhibitors were tested for their effect on elongation and wound healing when added to CTRL (purple) or CCM (orange). Taken together (A), the data present a non-linear relationship (blue curve) of phenotypic transformation between non-inhibitor conditions (white stars) and each inhibitor. The effects for each inhibitor were assessed by comparing each media condition with and without inhibitor (B-C). † *significant difference in at least one metric, * p* < *0*.*05, ** p* < *0*.*01, *** p* < *0*.*001, NS = no significant difference*

The data present a non-linear EC phenotype of wound healing and morphology (Figure 5A). The shape of the curve (blue)—determined by locally estimated scatterplot smoothing (LOESS)—suggests that as EC phenotype shifts from healthy CTRL (purple with white asterisk[*]) towards dysfunctional CCM (orange with white asterisk[*]), morphology and wound healing deteriorate together to a peak. Thereafter, elongation continues, and wound healing partially reverts albeit in less ordered form.

Overall, Akt and Ptp1b inhibition caused ECs in CCM to revert phenotype towards CTRL, suggesting increased activity in CCM. Inhibition of Src and Pdhk1, on the other hand, drove ECs in CTRL towards CCM dysfunction, suggesting decreased activity in CCM. The PTP1B/SHP2 inhibitor had no significant effect (Figure 5B-C).

Inhibition of Bmx in CCM decreased elongation and increased scratch area, matching the behavior of reverting from CCM to CTRL along the non-linear continuum. However, inhibition in CTRL increased elongation suggesting a worsening of phenotype from CTRL to CCM. Overall, the reversion from CCM was significantly larger (2.9-fold in elongation and 3.3-fold in scratch area).

## Discussion

The intact endothelial cell (EC) state and phenotype create a robust environment for optimized tissue repair. Quiescent ECs have maximal regulatory capacity (Nugent *et al*, 1993; Ettenson *et al*, 2000); whereas proliferative ECs (such as those involved in angiogenesis) can have opposite, dysfunctional and cancer promoting effects (Ferraro *et al*, 2019; Butler *et al*, 2010; Ghiabi *et al*, 2015). For years these precepts have guided interest in enhancing the retention and activity of anti-cancer drugs through regulation of tumor EC and vasculature, and in controlling tumor growth through regulation of angiogenic pathways (Jain, 2008; Giuliano & Pagès, 2013; Martin *et al*, 2019). We pursued further elucidation of the relationship between cellular phenotype and EC transformation in cancer and its molecular drivers and focused in particular on understanding the fine control that is achieved through the balanced activity of kinases and phosphatases in tumor vasculature

The focus on phosphoenzymes provides two advantages: their activity is well defined by the change in target phosphorylation, and they make excellent therapeutic targets. These enzymes act rapidly on cascading pathways, and a single phosphoenzyme often regulates several pathways (Schwartz & Murray, 2011). In anti-cancer therapy, protein kinases are now the major class of targets (Zhang *et al*, 2009). Furthermore, kinase inhibitors are the current state of the art for targeting tumor vasculature, with eleven recently approved or in phase II/III clinical trials (Qin *et al*, 2019). These target growth factor receptor (GFR) kinases, including VEGFR, PDGFR, c-Kit, and c-Met, primarily identified for their ability to revert vascular permeability and growth in angiogenic ECs, but to our knowledge have not been tested relative to initial EC state. Our study sought to identify the phosphoenzymes that drive the observed dysfunctional transformation in the confluent EC model.

Two functional metrics robustly separate the phenotypes of ECs in CCM from those in control media (CTRL): morphology (increased elongation) and wound healing (impaired response to monolayer injury) (Figure 1). To map the functional transformations to their phosphoenzymatic drivers, we collected quantitative phosphoproteomic data and validate for functional phenotype (Figure 2, 3); identified associated phosphoenzymes from KinomeXplorer predictions (Horn *et al*, 2014) (Figure 4); and probed with pharmacologic inhibitors against phosphoenzymes of interest (Figure 5). Phosphorylation state alone cannot directly predict protein activity in a high-throughput manner because of limited phosphosite annotation; instead, we used downstream phosphorylation as a measure of activity in upstream kinases and phosphatases.

Paradoxically, the aforementioned angiogenic GFR kinases all exhibited decreased activity in CCM, despite very elevated VEGF levels in the conditioned media, further reinforcing the need to explore mechanisms other than GF-driven angiogenesis. Indeed, though angiogenesis and the associated growth factors (GF) are important elements of cancer dynamics, they cannot account for the entirety of EC-cancer dynamics. For example, conditioned media from breast cancer cells (CCM) induces morphologic changes in confluent, quiescent human umbilical vein ECs in culture in an inversely proportionate manner with VEGF levels (Gomez *et al*, 2016). This suggests that there are important means of EC regulation in tumors in addition to classic GF-mediated angiogenesis. Hence, we focused on kinase and phosphatase families and selected six with the most differential significantly regulated activity in our analysis. These are all well-established players in tumor endothelial biology, and include: PDHK and AKT (predicted increased activity in CCM); SRC and TEC (decreased activity); and PTPN1/2 and PTPN11 (increased activity).

When pharmacologic inhibitors against these families were added to our culture model and assayed for morphology and wound healing, a non-linear pattern of EC phenotype along the two functional phenotype axes emerged (Figure 5A). The shape of the relationship suggests that while elongation increases linearly as EC phenotype shifts from CTRL to CCM, wound healing worsens beyond the CCM impairment, before partially recovering, albeit with dysmorphic EC and likely dysfunctional endothelium. Scratch closure is dependent on proliferation and migration: it may be that CCM dysfunction reduces proliferation while increasing migration, which is often associated with increased elongation (Dejana *et al*, 2018). The peak, then, represents the point at which migration overcomes decreased proliferation but does not imply restoration of a normal monolayer. The lower inflection of the curve is the result of remaining scratch area bounded on the by zero. A larger initial scratch and/or analysis with shorter time points would identify the continuum shape beyond CTRL. Further investigation with more inhibitors (and varying doses) is required to fully elucidate the underlying shape of the EC continuum along these axes.

The activity of each phosphoenzyme in CCM was determined from the effects of pharmacologic inhibition, compared to media without inhibitors (Figure 5B-C). Inhibition causing phenotypic reversion from CCM towards CTRL suggests *increased* activity of that phosphoenzyme in CCM. Conversely, an inhibitor driving dysfunctional phenotype from CTRL towards CCM suggests *decreased* activity in CCM. In this way, phosphoenzyme activity as predicted by phosphoproteomics can be compared to that determined by inhibition testing.

Both phosphoproteomics and inhibition testing confirmed increased activity of Akt in CCM, consistent with the increased Akt activity reported in angiogenic tumor ECs, and associated with elongated morphology (Guerrouahen *et al*, 2014; Tu *et al*, 2009; Cheng *et al*, 2017). However, given the observed decrease in angiogenic GFR kinase activity, the cancerous transformation in our model may follow an alternate Akt-based pathway that requires further investigation. As such, Akt kinase inhibitors are an interesting candidate for targeting both neo-angiogenesis and vessel co-option in cancer. They have been tested as anti-cancer agents in clinical trials including in metastatic TNBC (Nitulescu *et al*, 2016; Costa *et al*, 2018), but have not yet been tested for EC-regulating therapy.

Src activity is reported as a critical regulator of cytoskeletal structure and function including tumor EC permeability and angiogenesis (Indolfi *et al*, 2012; Sefton *et al*, 1981; Tinsley *et al*, 2013). In our confluent EC model, Src activity was *decreased* in both phosphoproteomics and inhibition testing. Decreased Src activity tracks with decreased activity of GFR kinases—which are upstream of Src—and overall, suggests that while Src inhibition may have regulating effects on angiogenic ECs, it may deepen the dysfunction of quiescent ECs. Such thought is further supported by the prominence of vascular toxicity in clinical trials using Src inhibitors as anti-cancer agents (Guignabert *et al*, 2016; Fujioka *et al*, 2018). Vascular adverse events have also been seen in anti-angiogenesis trials with apatinib, which targets Src kinases as well as VEGFR-2 and c-Kit (Qin *et al*, 2019).

Similar to Src, the phosphatase Ptp1b (PTPN1) is reported to show decreased activity in angiogenic ECs, since Ptp1b is a negative regulator of VEGF (Lanahan *et al*, 2014), and our model system stayed true to form. Transformation of quiescent ECs in our model showed *increased* activity in CCM, and as with Src, this apparent discrepancy is likely linked to decreased GFR kinase activity. Therefore, while our data suggest Ptp1b as a candidate target for quiescent EC regulation by inhibition, further investigation is required for angiogenic ECs.

Pdhk1 (the most represented PDHK family kinase in our dataset) is involved in metabolic regulation (Zhang *et al*, 2016). Pharmacologic inhibition of Pdhk1 in ECs in CTRL drove CCM dysfunction, consistent with previously reported suppression of EC proliferation (Huang *et al*, 2015). Pdhk1 inhibition has been suggested as an EC-regulating and anti-cancer therapy (Huang *et al*, 2015; Sutendra & Michelakis, 2013), but these data suggest caution. Furthermore, inhibition in CCM had no significant effect, suggesting that Pdhk1 activity is already very low in CCM. Notably, the pharmacologic data for Pdhk1 are opposite to that predicted by phosphoproteomics and KinomeXplorer, highlighting limitations of our method: accurate estimates require broad coverage of the phosphosites targeted by the phosphoenzyme of interest; in our study, only three sites had significant predicted interactions with Pdhk1. The analysis, thereby, depends on the balance between increased accuracy and decreased coverage as a result of more stringent NetworKIN score thresholding.

Bmx (of the TEC family, also known as Etk, the endothelial and epithelial tyrosine kinase) is an important regulator of EC cytoskeletal reorganization (Cenni *et al*, 2012). It is cross-activated by VEGFR and Src, and plays an important in VEGF-induced angiogenesis (Zhang *et al*, 2003). Consistent with this, phosphoproteomic analysis predicted reduced Bmx activity in CCM, consistent with reduced activity of angiogenic GFR kinases and Src. However, pharmacologic inhibition of Bmx paradoxically caused a large reversion from CCM (suggesting increased activity in CCM) and a smaller induction of dysfunction from CTRL (decreased activity). The paradoxical result may indicate that Bmx dysfunction in cancer occurs via a combination of pathways, angiogenic and non-angiogenic, and may also reflect a phenotype-dependent response. Bmx kinase inhibitors have been considered as potential EC-regulating therapies in cancer (Jarboe *et al*, 2013), but our results strongly suggest further investigation is required.

Finally, the broad-spectrum Ptp1b/Shp2 inhibitor showed no significant effect. This could be caused by the limited strength of the inhibitor (maximal non-cytotoxic dose of 40 µM was only 20-40x IC50). It could also reflect the broad spectrum of phosphatases targeted by the inhibitor, some of which induce dysfunction while others inhibit it. While KinomeXplorer analysis of phosphatases predicted increased activity for all phosphatases, there are both pro- and anti-tumorigenic phosphatases (Labbé *et al*, 2012). This artifact is likely due to the limited coverage of phosphatases in KinomeXplorer which only covers a subset of the protein tyrosine phosphatase (PTP) family. Vascular endothelial protein tyrosine phosphatase (VE-PTP), for example, is not represented, but it is known to play a critical role in regulating EC cytoskeletal changes and has been considered as a target for EC regulation in cancer (Kontos & Willett, 2013).

Altogether, quiescent, confluent ECs in CCM show increased Akt activity, but downregulation of GFR kinase pathways. This behavior is significantly different from that reported for angiogenic ECs, and suggests additional mechanisms of EC-cancer dysfunction in quiescent, healthy vessels co-opted by tumors compared to neo-angiogenic tumor vessels. Our data support Akt inhibition for normalizing EC phenotype, and advocate for caution in inhibition of angiogenic targets—such as VEGFR and Src— that may have adverse effects on healthy blood vessels.

Overall it is fascinating to see that the integrated effects of rapidly acting co-regulatory kinases and phosphatases can establish a dynamic of great potency and dimensionality. Phosphoproteomics offers news insights into tumor EC biology and possibly a new test bed for emerging chemotherapies. We can now offer yet another aspect of the complex continuum that is the tumor microenvironment and sophistication of EC biology and phenotype spectrum.

## Materials and Methods

### Endothelial and breast cancer cell culture

Primary human umbilical vein endothelial cells (HUVEC, Lonza) were used in passages 3-6 and cultured on 0.1% gelatin-coated tissue culture plates in complete EGM2 (Lonza) supplemented to a total of 10% Fetal Bovine Serum (FBS). Cells were passaged by detachment with trypsin with removal of debris by centrifugation (5 mins, 300g), and reseeded at 10k cells/cm^2^. Cells were cultured under standard conditions (37°C, 5% CO_2_). Human MDA-MB-231 breast cancer cells (ATCC) were cultured on tissue culture plates in Dulbecco’s Modified Eagle Medium (DMEM) with L-Glutamine (Gibco) supplemented with 10% FBS and 1% penicillin/streptomycin. Cells were cultured under standard conditions and expanded and stored according to the manufacturer’s recommendations.

### Breast cancer conditioned media

To account for reduced bicarbonate levels in EGM2, we instead used RPMI 1640 (Gibco) as the base of the CTRL media, which was supplemented with 10% FBS and the EGM2 bullet kit containing VEGF, rEGF, rbFGF, IGF, heparin, hydrocortisone, ascorbic acid, and GA-1000 (antibiotic). To ensure sufficient bicarbonate levels, the media was supplemented with sodium bicarbonate to a final concentration of 3.7g/L. MDA-MB-231 cells were seeded at 10k cells/cm^2^ and grown for 4-6 days. Media was removed and the cells were washed one time with PBS. The cells were then grown in fresh CTRL media for 2 days before collection. Debris and cells were removed by centrifugation for 15 mins at 1,000g. To account for glucose depletion, 2 g/L glucose was added; however, experiments were also successful without supplementation. For consistency, this cancer-conditioned media (CCM) was always stored at -80°C for at least two days before use in EC cultures.

### Endothelial culture in cancer-conditioned media (CCM)

HUVECs were seeded at 10k cells/cm^2^ and grown to confluence (approximately 4 days). After one quick wash (∼15s) with PBS containing Ca^2+^ and Mg^2+^ to wash without detachment, either 100% CTRL (described above) or 75% CCM + 25% CTRL was added, with and without inhibitors.

### Kinase and Phosphatase Inhibitors

The following six inhibitors were used in this study from Calbiochem: PDHK inhibitor, AZD7545; AKT inhibitor, AT7867; BMX inhibitor, QL47; SRC inhibitor, PP2; PTP1B (also known as PTPN1) inhibitor, JTT-551; and HePTP inhibitor inhibitor, ML119 (which targets several phosphatases including PTPN1 and PTPN11 (Expanded View Figure 3).

### Phase contrast and fluorescence microscopy

EC cultures were imaged throughout culture using phase contrast microscopy. At the end of the experiment, EC cultures were fixed for fluorescent staining and analysis, by addition of paraformaldehyde (PFA) directly to the cultures for a final concentration of 2-4% PFA, without washing in order to prevent potential alteration of morphology during processing. Cells were fixed for 10-15 mins at room temperature, and then permeabilized with 0.3% Triton X-100. Cells and nuclei were stained using Cell Tracker Orange CMTMR (Thermo) at 10 µM and DAPI (Thermo) at 0.5 µM, respectively, for 30 minutes at room temperature. Microscopy was performed using a Nikon Ti-E microscope with 4X, 10X, or 20X magnification, a Prior ProScan III motorized stage, and NIS Elements AR acquisition software. Images were captured using a Photometrics CoolSnap EZ or an Andor Zyla 4.2 camera and analyzed using NIS Elements or ImageJ.

### Endothelial morphology in CCM

Tiled fluorescent images of ECs stained with Cell Tracker Orange CMTMR, as described above were acquired at 10X magnification for a final square image of side length 4.93 mm (7578 px). To reduce boundary effects, a 3000 × 3000 px area in the middle was cropped for analysis. ImageJ was used for automatic segmentation and quantification of cell shape by the following steps: automatic contrast enhancement, smoothing; automatic thresholding by mean brightness; filling holes; smoothing edges and rebinarizing; and measuring using “Analyze Particles…”, filtering objects smaller than 1000 and larger than 5000 pixels in size, and well as excluding objects touching the edge. Subsequent quantitative analysis was done using the R programming language.

### Endothelial response to scratch injury

ECs were grown until confluent as described above. A controlled cross scratch injury was then made using a p1000 pipette tip (in a single vertical motion followed by a single horizontal motion). The media was then removed and replaced with CTRL or CCM, with or without inhibitors. The scratch area was tracked by imaging at 0h, 16h, 25h, and 36h using phase contrast microscopy—5×5 tiled images taken at 4X magnification for a total image area of 9.3 × 9.3 mm. Scratch area was quantified by manual segmentation on ImageJ analysis software.

### Phosphoproteomic sample preparation

Cells were lysed in 8M urea (Sigma) with protease and phosphatase inhibitors (Thermo) and quantified using BCA assay kit (Pierce). Proteins were reduced with 10mM dithiothreitol (Sigma) for 1h at 56°C and then alkylated with 55mM iodoacetamide (Sigma) for 1h at 25°C in the dark. Proteins were then digested with modified trypsin (Promega) at an enzyme/substrate ratio of 1:50 in 100mM ammonium acetate, pH 8.9 at 25°C overnight. Trypsin activity was halted by addition of acetic acid (99.9%, Sigma) to a final concentration of 5%. After desalting using Protea C18 spin columns, peptides were lyophilized in 400ug aliquots and stored at -80°C.

### Peptide labeling, enrichment, and fractionation

Peptide labeling with TMT 10plex (Thermo) was performed per manufacturer’s instructions. Lyophilized samples were dissolved in 70 μL ethanol and 30 μL of 500 mM triethylammonium bicarbonate, pH 8.5, and the TMT reagent was dissolved in 30 μL of anhydrous acetonitrile. The solution containing peptides and TMT reagent was vortexed and incubated at room temperature for 1 h. Samples labeled with the eight different isotopic TMT reagents were combined and concentrated to completion in a vacuum centrifuge. For immunoprecipitation, protein G agarose (60µL, Millipore) was incubated with anti-phosphotyrosine antibodies (12µg 4G10 (Millipore), 12µg PT66 (Sigma), and 12µg PY100 (CST)) in 400µL of IP buffer (100mM Tris, 100mM NaCl, and 1% Nonidet P-40, pH 7.4) for 6h at 4°C with rotation. The antibody conjugated protein G was washed with 400µL of IP buffer. The TMT labeled peptides were dissolved in 400µL IP buffer and the pH was adjusted to 7.4. The TMT labeled peptides were then incubated with the antibody conjugated protein G overnight at 4°C with rotation. The supernatant from the IP was collected and stored at -20C. The agarose was washed with 400µL IP buffer followed by four rinses with 400µL rinse buffer (100mM Tris, pH 7.4). Peptides were eluted with 50µL of 0.2% TFA for 30 minutes at 25°C. Immobilized metal affinity chromatography (IMAC) was used to further enrich for phosphopeptides using the High-Select Fe-NTA phosphopeptide enrichment kit (Thermo) per manufacturer’s instructions. Elution was analyzed via LC-MS/MS as described below. The supernatant from the pY IP was fractioned via high-pH reverse phase HPLC. Peptides were resuspended in 100uL buffer A (10mM TEAB, pH8) and separated on a 4.6mm x 250 mm 300Extend-C18, 5um column (Agilent) using an 90 minute gradient with buffer B (90% MeCN, 10mM TEAB, pH8) at a flow rate of 1ml/min. The gradient was as follows: 1-5% B (0-10min), 5-35% B (10-70min), 35-70% B (70-80min), 70% B (80-90min). Fractions were collected over 75 minutes at 1 minute intervals from 10 min to 85 min. The fractions were concatenated into 10 fractions non-contiguously (1+11+21+31+41+51+61+71, 2+12+22+32+42+52+62+72, etc). The fractions were concentrated in a vacuum centrifuge and then were lyophilized. Phosphopeptides were enriched from each of the 15 fractions using the High-Select Fe-NTA phosphopeptide enrichment kit (Thermo) per manufacturer’s instructions. Each elution was analyzed via LC-MS/MS as described below. The supernatant from each fraction was collected and stored at -20C. For protein expression, each supernatant was diluted and analyzed via LC-MS/MS as described below.

### LC-MS/MS

Peptides were loaded on a precolumn and separated by reverse phase HPLC using an EASY-nLC1000 (Thermo) over a 140 minute gradient before nanoelectrospray using a QExactive mass spectrometer (Thermo). The mass spectrometer was operated in a data-dependent mode. The parameters for the full scan MS were: resolution of 70,000 across 350-2000 *m/z*, AGC 3e^6^, and maximum IT 50 ms. The full MS scan was followed by MS/MS for the top 10 precursor ions in each cycle with a NCE of 32 and dynamic exclusion of 30 s. Raw mass spectral data files (.raw) were searched using Proteome Discoverer (Thermo) and Mascot version 2.4.1 (Matrix Science). Mascot search parameters were: 10 ppm mass tolerance for precursor ions; 15 mmu for fragment ion mass tolerance; 2 missed cleavages of trypsin; fixed modification were carbamidomethylation of cysteine and TMT 10plex modification of lysines and peptide N-termini; variable modifications were methionine oxidation, tyrosine phosphorylation, and serine/threonine phosphorylation. TMT quantification was obtained using Proteome Discoverer and isotopically corrected per manufacturer’s instructions, and were normalized to the mean of each TMT channel. Only peptides with a Mascot score greater than or equal to 25 and an isolation interference less than or equal to 30 were included in the data analysis.

### Phosphoproteomic Data Post-Processing

Any Ser or Thr site assignments among pTyr-enriched fractions were flagged and corrected to a neighboring Tyr within the same peptide. If multiple Tyr were present in the sequence, a Tyr site was chosen that had already been registered by another peptide with equivalent sequence, or, alternatively, the one that was best supported by LTP/HTP references from PhosphoSitePlus (Hornbeck *et al*, 2012). The data was then merged by summing raw LC-MS/MS readout. Occasionally, peptides were assigned to multiple protein sources, but some methods of analysis required single protein assignments. Most conflicts were resolved by choosing recognized proteins over putative proteins, while occasionally a protein was chosen because it was better researched, either for the specific phosphosite (by PhosphoSitePlus) or in general.

Proteins were classified as cytoskeletal if their Gene Ontology (GO) collection annotations (specifically from GO Biological Processes and GO Cellular Component (Ashburner *et al*, 2000; The Gene Ontology Consortium, 2019)) matched with one of the following cytoskeletal keywords (* = wild card): *actin, adhesion, cytoskelet*, edge, junction, microtubule, *podium, projection*.

### Western blot

EC cultures were lysed on ice in RIPA buffer (Sigma) with protease and phosphatase inhibitors (Thermo). A cell scraper was then used to collect the partial lysate and lysing was then continued at 4°C for 30 minutes. The insoluble fraction was removed by centrifugation spun at 16,000g for 30 minutes at 4°C, and the protein content quantified using BCA Protein Assay Kit (Pierce) according to the manufacturer’s instructions. Lysate was prepared with 4x LDS Buffer and 10x Reducing Agent (Life Technologies) according to the manufacturer’s instructions, denatured at 95°C for 5 minutes, and stored at -20°C. Thawed samples were run on pre-cast 4-12% or 10% Bis-Tris gels (Life Technologies) using NuPage MOPS Buffer (Life Technologies), in an XCell SureLock Mini-Cell electrophoresis system (Invitrogen). Membrane transfer was completed using the iBlot transfer system according to the manufacturer’s instructions. Membranes were blocked with 5% non-fat dry milk for 1 hour at room temperature. All antibodies were acquired from Cell Signaling Technologies (#4539, #4191, #89930, #5651, #3873), and diluted in 10% Starting Block buffer (Thermo). Primary antibodies were incubated overnight at 4°C. HRP-conjugated secondary antibodies (Invitrogen) were incubated for 1 hour at room temperature. Chemiluminescent detection was done using Immobilon Forte Western HRP Substrate (Millipore) and a ChemiDoc XRS+ Imaging system (BioRad).

### Data Analysis and Statistics

All statistical comparisons were done using the Student’s t-test, unless otherwise indicated, and a Benjamini-Hochberg false discovery correction was then applied for multiple comparisons. All data analysis was completed using the R programming language (R Core Team, 2018) and the following packages: *Tidyverse* package for data processing (Wickham, 2017); *Stats* core R package for nonlinear least squares (*nls*) and hierarchical clustering (*hclust*); *sigclust2* package for cluster separation analysis (Kimes *et al*, 2017); and *ggplot2* (Wickham, 2016), *ggpubr* (Kassambara, 2018), and *pheatmap* (Kolde, 2019) for visualizations.

### Data Availability

The mass spectrometry proteomics data have been deposited to the ProteomeXchange Consortium via the PRIDE (Perez-Riverol *et al*, 2019) partner repository with the dataset identifier PXD020333 and 10.6019/PXD020333.

## Acknowledgements

The authors wish to thank Dr. Amanda del Rosario, Dr. Antonius Koller, and the Koch Institute Proteomics Core for assistance with quantitative mass spectrometry and downstream analysis. This work was supported in part by an HST Martinos Research Scholarship and a Hugh Hampton Young fellowship to O.G., and a NIH grant R01 49039 to O.G. and E.R.E.

## Author contributions

Conceptualization, O.G. and E.R.E.; Methodology, O.G. and E.R.E.; Formal Analysis, O.G.; Investigation, O.G.; Data Curation, O.G.; Writing – Original Draft, O.G. and E.R.E.; Writing – Review & Editing, O.G. and E.R.E.; Visualization, O.G.; Supervision, E.R.E; Funding Acquisition, E.R.E.

## Conflict of Interest

The authors declare no conflict of interest.

## Expanded View Figures

**Expanded View Figure 1.**
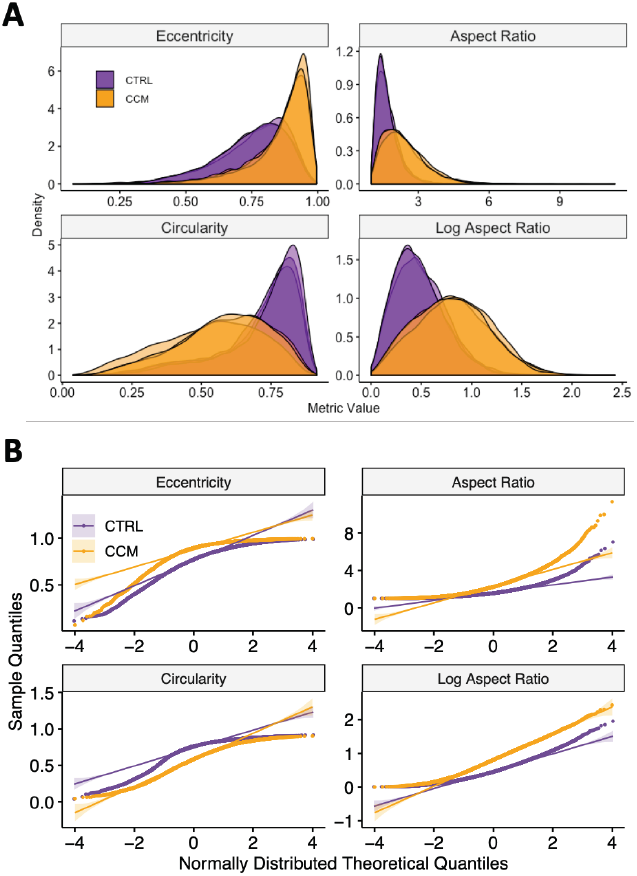
Log Aspect Ratio is the best metric to quantify the elongation phenotype between ECs in CTRL and CCM. HUVECs show variable elongation after 36 hours in control (CTRL, purple) or cancer-conditioned media (CCM, orange). Kernel-estimated density plots of four metrics of elongation across three replicates yield the best separation between phenotypes using Log Aspect Ratio (A). Quantile-quantile plots show that the Log Aspect Ratio is the most normally distributed.

**Expanded View Figure 2.**
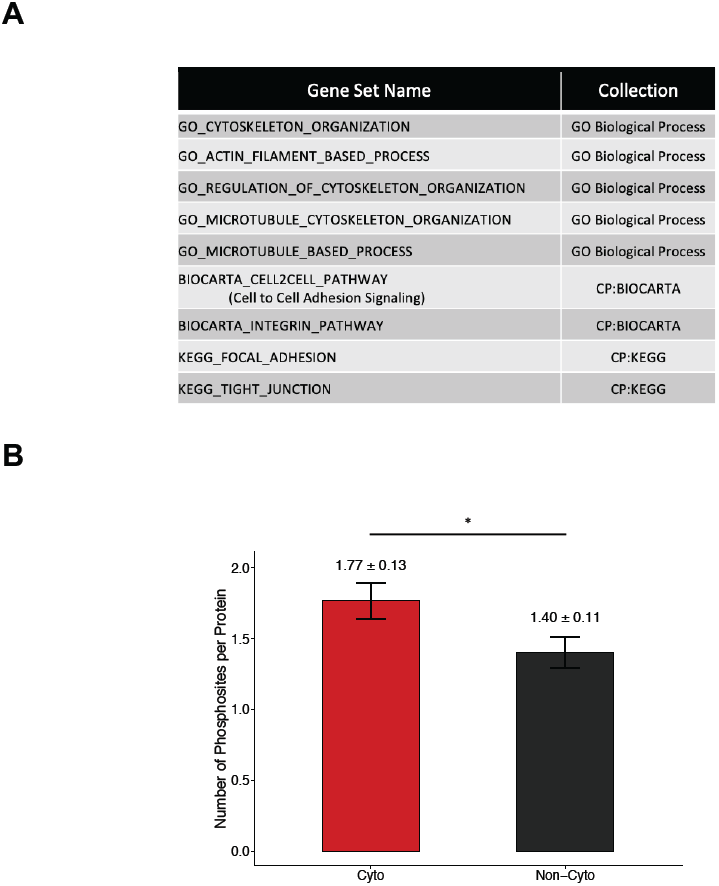
Phosphoproteomic data is further associated with cytoskeletal changes. Import cytoskeletal gene sets were found significantly enriched among the phosphoproteomic data by gene set overlap analysis with Gene Ontology (GO) Biological Process (Ashburner *et al*, 2000; The Gene Ontology Consortium, 2019), BioCarta (Nishimura, 2002) and KEGG (Kanehisa & Goto, 2000; Kanehisa *et al*, 2017, 2019) (A). Cytoskeletal proteins had significantly higher phosphorylations per protein than non-cytoskeletal proteins (B). *\* p* < *0*.*05*

**Expanded View Figure 3.**
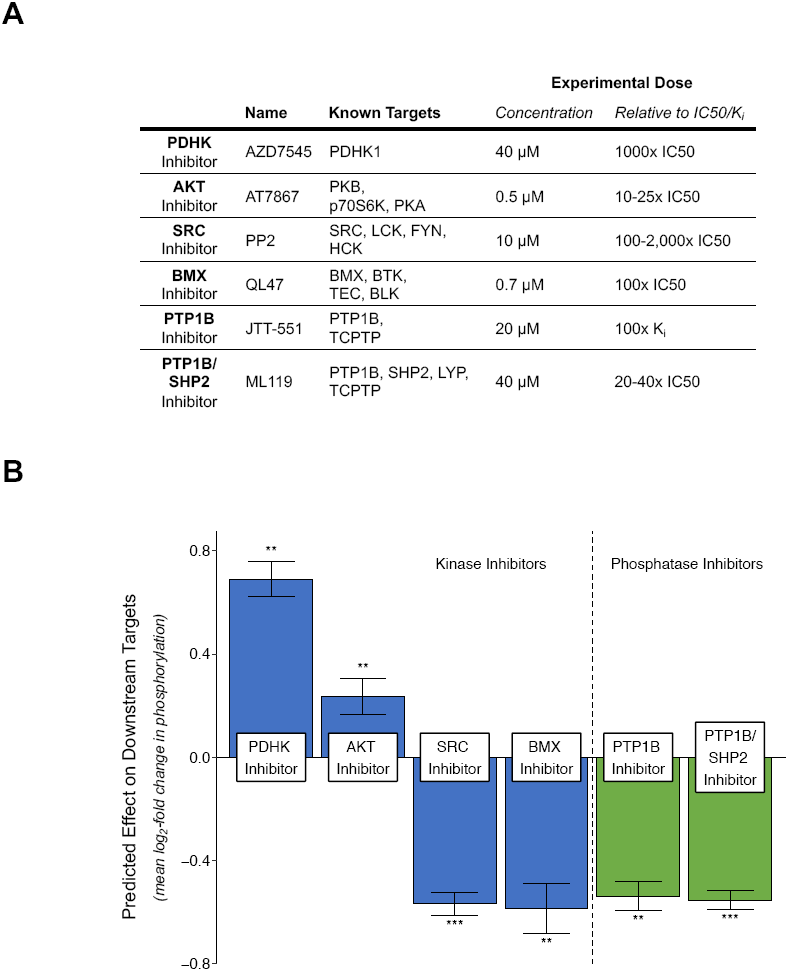
Secondary targets of chosen pharmacologic inhibitors do not affect predicted activity by KinomeXplorer analysis. Six pharmacologic inhibitors were chosen against four kinase (blue) and two phosphatase (green) families with significant predicted changes in activity between CTRL and CCM by KinomeXplorer analysis (A). Repeated predictive analysis taking into account each inhibitor’s targets—including secondary ones—did not significantly affect the predictions (B). *\* p* < *0*.*05, ** p* < *0*.*01, *** p* < *0*.*001*

